# Parameter estimation for correlated Ornstein-Uhlenbeck time-series

**DOI:** 10.1101/2021.02.12.430978

**Authors:** Helmut H. Strey, Rajat Kumar, Lilianne Mujica-Parodi

## Abstract

In this article, we develop a Maximum likelihood (ML) approach to estimate parameters from correlated time traces that originate from coupled Ornstein-Uhlenbeck processes. The most common technique to characterize the correlation between time-series is to calculate the Pearson correlation coefficient. Here we show that for time series with memory (or a characteristic relaxation time), our method gives more reliable results, but also results in coupling coefficients and their uncertainties given the data. We investigate how these uncertainties depend on the number of samples, the relaxation times and sampling time. To validate our analytic results, we performed simulations over a wide range of correlation coefficients both using our maximum likelihood solutions and Markov-Chain Monte-Carlo (MCMC) simulations. We found that both ML and MCMC result in the same parameter estimations. We also found that when analyzing the same data, the ML and MCMC uncertainties are strongly correlated, while ML underestimates the uncertainties by a factor of 1.5 to 3 over a large range of parameters. For large datasets, we can therfore use the less computationally expensive maximum likelihood method to run over the whole dataset, and then we can use MCMC on a few samples to determine the factor by which the ML method underestimates the uncertainties. To illustrate the application of our method, we apply it to time series of brain activation using fMRI measurements of the human default mode network. We show that our method significantly improves the interpretation of multi-subject measurements of correlations between brain regions by providing parameter confidence intervals for individual measurements, which allows for distinguishing between the variance from differences between subjects from variance due to measurement error.

## A. Introduction

In many fields of science one is interested in whether two sets of data are linearly correlated. One of the simplest statistical tool for such a case is to calculate the Pearson correlation coefficient *ρ*. By calculating the Pearson correlation coefficient we assume that the samples are identically and independenly drawn from a bivariate Gaussian distribution. Unfortunately, the Pearson coefficient is frequently used for data that falls outside this condition. In particular, when applying Pearson correlations to time-series that may be distributed normally but where adjacent time points are not independent. In this article, we are developing a method to characterize correlations between two time-series that exhibit short-term memory or relaxation. In particluar, we will restrict ourselves to the simplest random process that results in a fluctuating time series with a characteristic relaxation time: the Ornstein-Uhlenbeck (OU) process [1]. A OU process with mean zero can be expressed by the following Langevin equation:

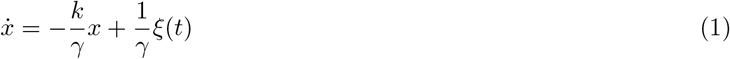

where k is the spring constant, γ is the friction coefficient and ξ(*t*) is a randomly fluctuating force. We and others have recently published maximum likelihood methods to estimate parameters from an OU process time series [2–4], and here we attempt the same for coupled OU processes.

## B. Coupled Oscillators

We consider a system of two overdamped oscillators, characterized by a friction coefficient γ and spring constant *k*, that are coupled by a spring with a spring constant *c*. The Langevin equations can be written in the following way:

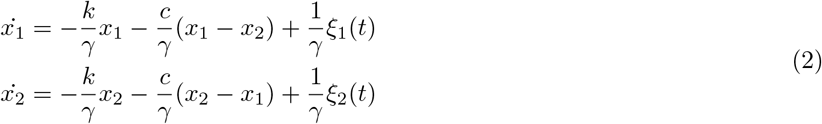

This system can be decoupled by the two eigenfunctions 2*y*_1_ = *x*_1_ + *x*_2_ and 2*y*_2_ = *x*_1_ *− x*_2_, which results in

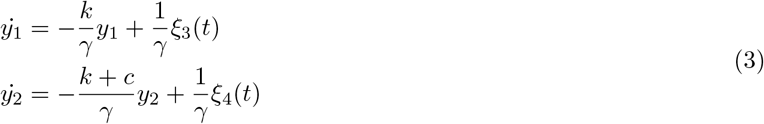

where *y*_1_(*t*) and *y*_2_(*t*) are two Ornstein-Uhlenbeck processes with different relaxation rates. The equipartition theorem yields 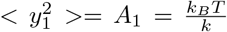 and 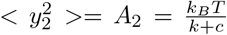. The other important relationships are: diffusion coefficient 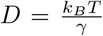, relaxation times 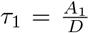 and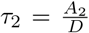. If the friction coefficients γ are different for *x*_1_ and *x*_2_ one can still decouple the system of differential equations by finding the eigenvectors and eigenvalues [5]. For simplicity we assume that both processes have the same friction coefficient or diffusion coefficient *D*.

In many fields, the Pearson Correlation Coefficient *ρ* is used to characterize the correlation between two time-series. The connection between our coupling coefficient *c* and *ρ* is:

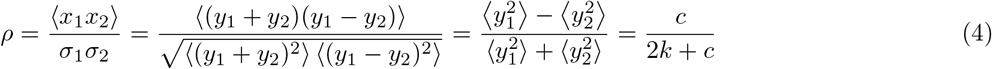

As expected, for *c* = 0 we obtain *ρ* = 0, and *c* = ∞ results in *ρ* = 1. Often correlation analysis are performed on normalized samples. It therefore makes sense to define a unitless coupling coefficient *C* = *c/k* which can be calculated by estimating *A*_1_ and *A*_2_ from the data:

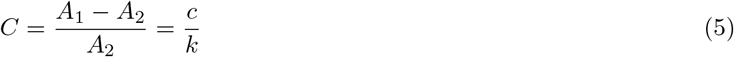

We can extend this procedure to negative coupling coefficients. For example, *c* = −*k* would results in *ρ* = −1, but at this *c* (see eq. 3) *y*_2_ becomes unbound. In order to map the normalized coupling coefficient C in the range from −∞ to +∞ to the Pearson correlation coefficient *ρ* = −1 to *ρ* = 1 we suggest the following procedure: From two normalized samples *x*_1_ and *x*_2_, we calculate 2*y*_1_ = *x*_1_−*x*_2_ and 2*y*_2_ = *x*_1_−*x*_2_. From *y*_1_ and *y*_2_, we estimate the amplitudes *A*_1_ and *A*_2_. If *A*_1_ *> A*_2_ then the coupling coefficient *C >* 0 and can be calculated using *C* = (*A*_1_− *A*_2_)*/A*_2_. If on the other hand *A*_1_ *< A*_2_ then the coupling coefficent is negative and *C* = (*A*_1_−*A*_2_)*/A*_1_. Reversing the process, we can use this procedure to create artificial time-series, by simulating two independent OU processes (eq. 3) to produce two coupled OU processes *x*_1_ = *y*_1_ + *y*_2_ and *x*_2_ = *y*_1_ *− y*_2_ with a well-defined positive coupling *C* or *ρ*. One obtains negative coupling coefficients by using *x*_2_ = *y*_2_ *− y*1.

## C. Maximum likelihood solution

In this section we will express the likelihood (see for example [6]) of two correlated time-series *x*_1_ and *x*_2_ in terms of *A*_1_, *A*_2_ and 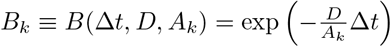assuming uniform priors for all parameters.

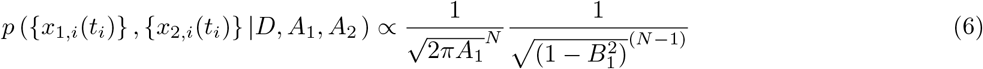

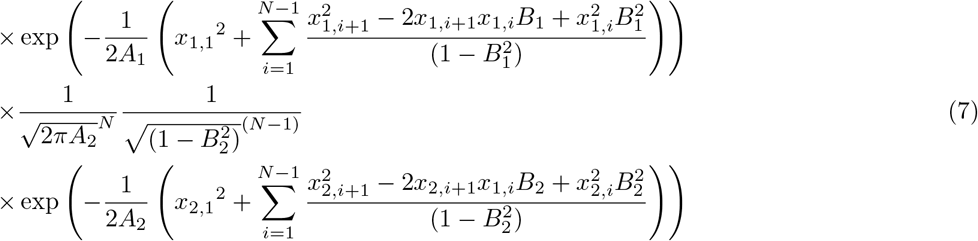

In order to find the maximum likelihood it is convenient to take the logarithm of *p* and then take the derivatives with respect to *A*_1_, *A*_2_ and *D*.

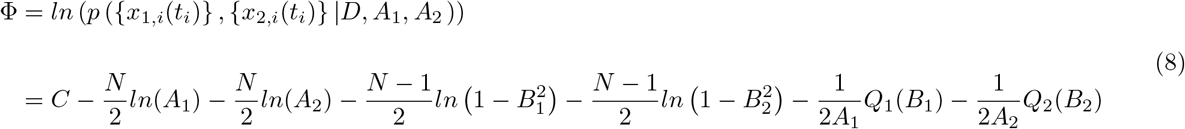

with

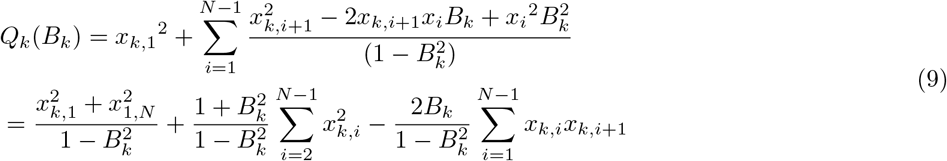

with *C* representing an unimportant constant. *Q_k_*(*B_k_*) reveals the fundamental statistic - the only terms that exclusively contain the data {*x_k,i_*} with *k* = 1, 2:

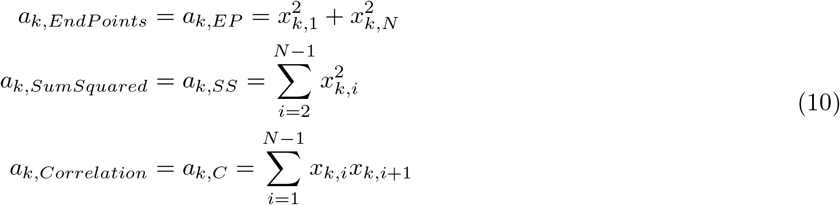

We find the maximum likelihood parameters *A_k,max_* and *D_max_* by finding the roots of the first derivatives with respect to the three parameters. The derivative with respect to *A_k_* results in the following two conditions:

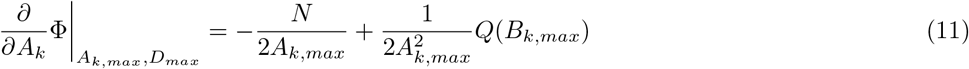

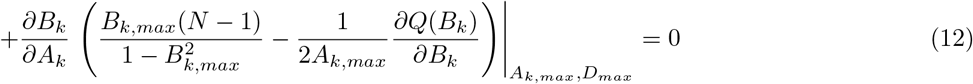

with

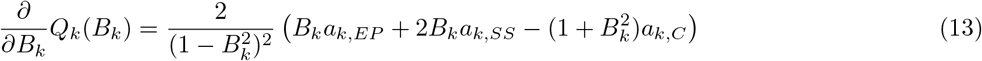

and

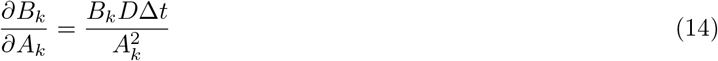

The derivative with respect to *D* can be written:

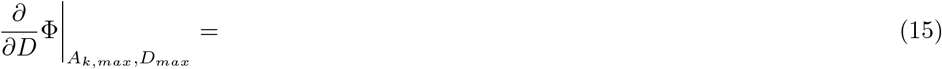

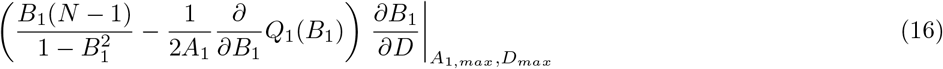

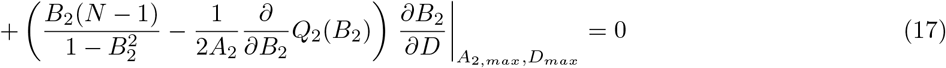

with

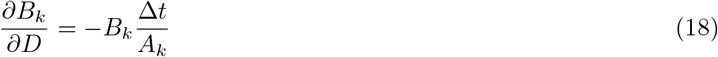

From these three equations we need to find *A*_1_,*A*_2_ and *D*, which can be done using numerical root finding algorithms that are implemented in standard numerical libraries (e.g. Powell hybrid method implemented in MINPACK [7]). In order to estimate the parameter uncertainties, we need to calculate the Jacobian. Here the only non-zero terms are:

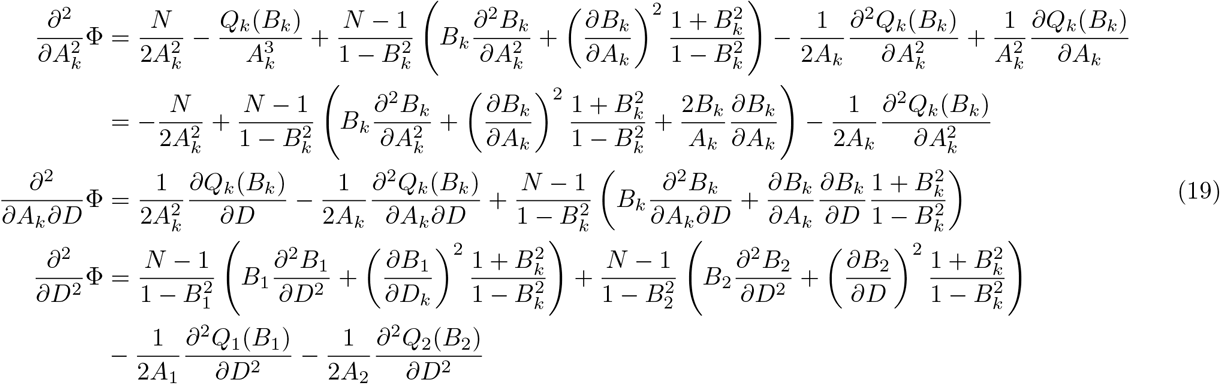

with

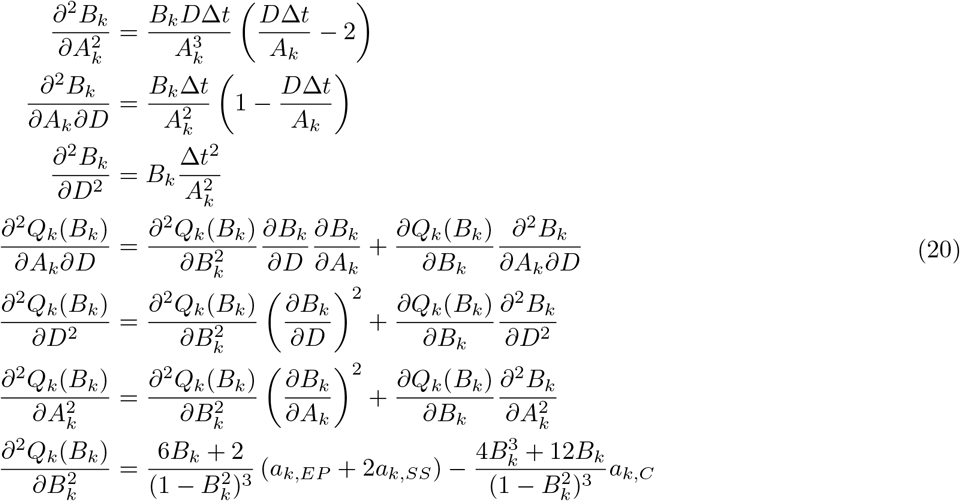

Using these terms, we can now express the Hessian *H* as:

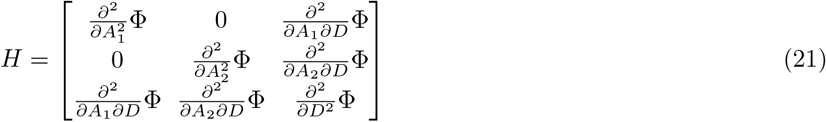

whose negative inverse determines the parameter’s covariance matrix *C* = − *H^−^*^1^(*A*_1*,max*_, *A*_2,max_, *D*_*max*_) (see for example [6]). In order to get a better sense of how the parameter estimation errors behave we can calculate the expectation values of the second derivaties considering *N* →∞: ⟨*a_k,EP_*⟩ = 2 *A_k_*, ⟨*a_k,SS_*⟩ = (*N* − 2)*A_k_*, and ⟨*a_k,C_*⟩ = (*N* − 1)*A_k_B_k_*. This results in the following expectation values of the derivatives of *Q_k_*(*B_k_*):

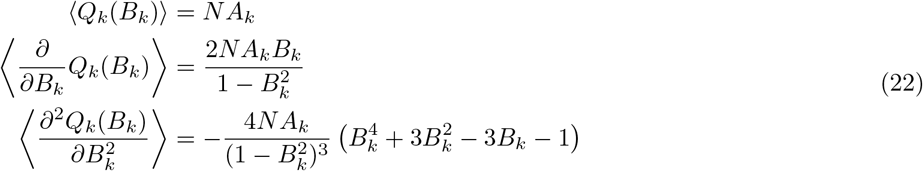

From this, we can express the Hessian using the following second derivatives:

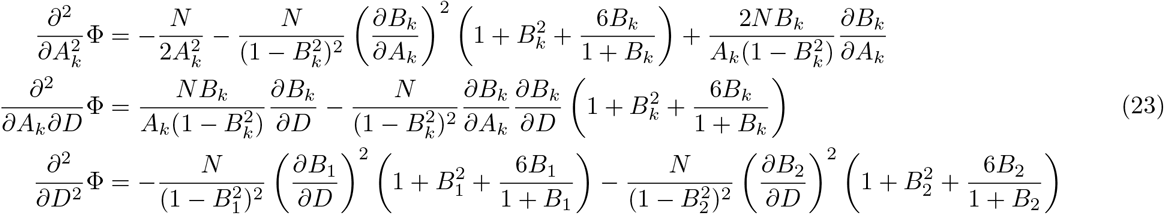

These equations allow us to express how the uncertainties/errors of the parameters depend on the sampling interval ∆*t/τ* given that we measure *N* samples. In Fig. 2 we plot the relative errors of *A*_1_*, A*_2_ and *C* as a function of ∆*t/τ*_1_ and *ρ* for large fixed *N* (*N* = 10^6^) using eqs. 23. For example, in Fig.2a we can see that the optimal sampling interval as given by ∆*t/τ*_1_ is reducing as *ρ* is increasing. This is most likely due to the fact that the relaxation *τ*_2_ is decreasing as *ρ* is increasing which will affect the uncertainty of both *A*_1_ and *A*_2_. We also observe a shift in behavior for larger *ρ >* 0.25 where no minimum ∆*t/τ*_1_ for *A*_1_ and *C* exists. This is in contrast to our finding for a single OU process [2] where the relaxation time *τ* had a minimum of approximatly ∆*t* = 0.7*τ* and *dA* continuously decreased as ∆*t/τ* decreased.

**FIG. 1:**
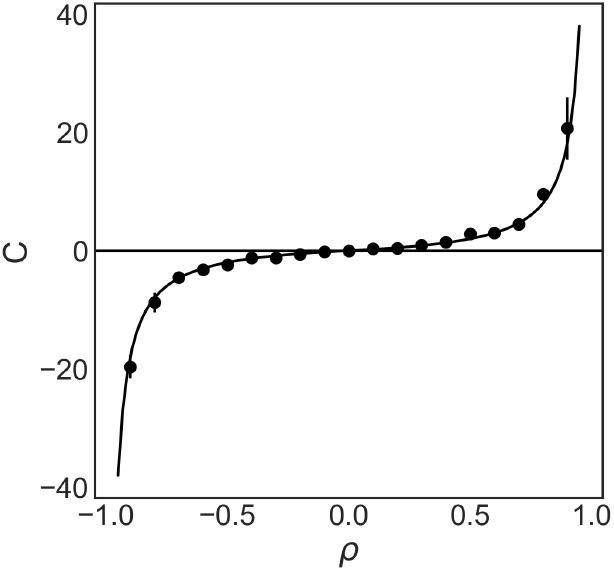
Relationship between coupling coefficient *C* and Pearson correlation coefficient *ρ*. The solid line represents eq. 5, whereas the data points represent the parameter estimation by MCMC simulation on artificial data of length *N* = 1000. The errorbars indicate the distribution of estimated *C*s.

**FIG. 2:**
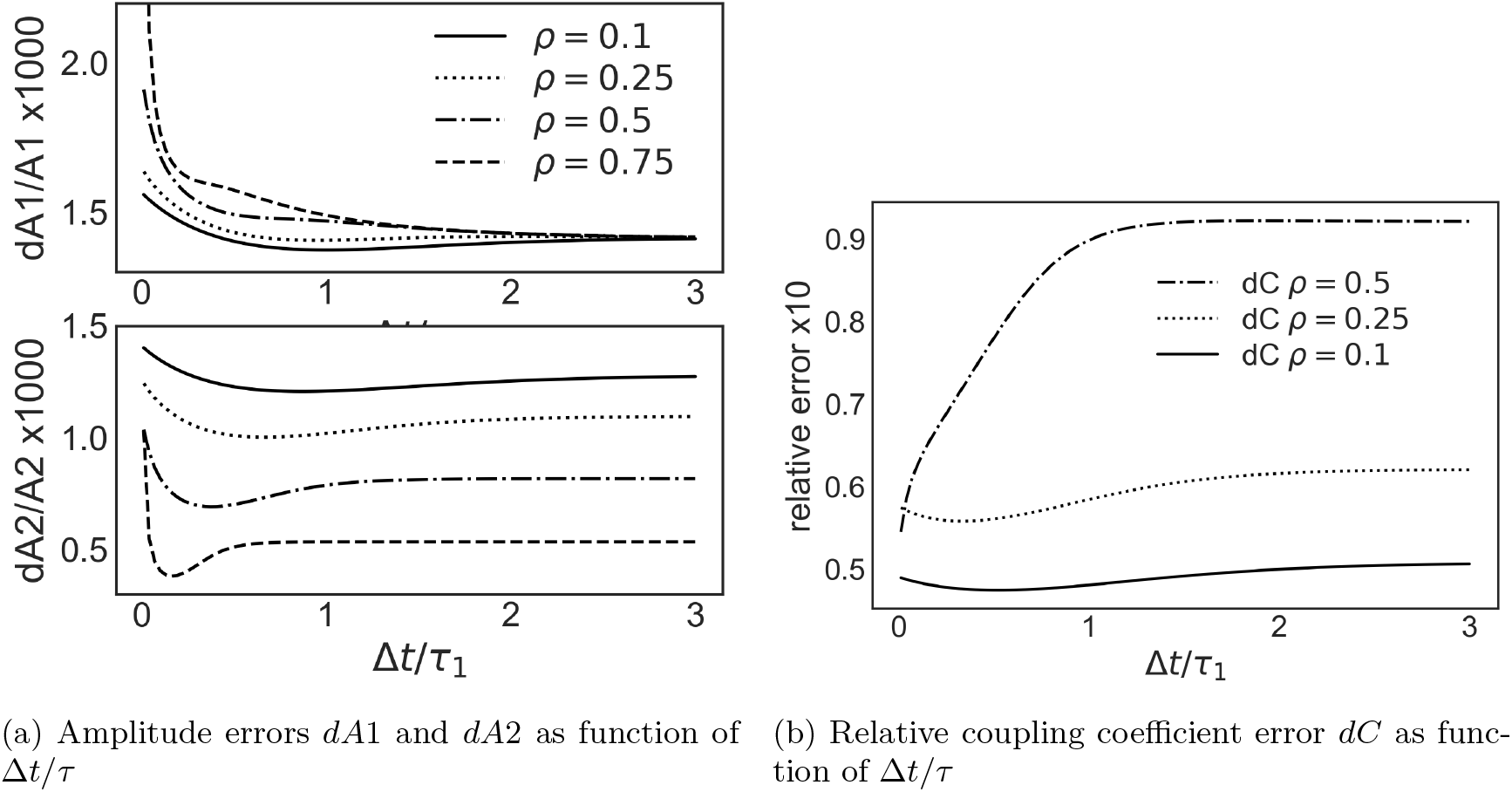
Dependence of relative errors on ∆*t/τ* for large *N* (*N* = 10^6^) using eqs. 23

## D. Validation by Simulation

In this section, we create artifical correlated time-series employing the method mentioned in B to validate our Maximum-Likelihood method against parameter estimation by Markov-Chain Monte-Carlo (MCMC) methods [8]. Probabilistic calculations using MCMC were carried out in python and Julia, using the anaconda distribution of python [9] with PYMC3 [10] and Julia 1.53 [11] with Turing.jl [12]. For the following table and figures, we created 400 correlated time-series pairs for Pearson Correlation coefficients of 0.9, 0.5, 0.25, and 0.1 for *N* = 1000 at *D* = 1*, A*_1_ = 1, ∆*t* = 0.3. Since the problem is symmetric with respect to the sign of the correlation coefficient we don’t need to simulate negative correlation coefficients. Fig. 3 shows the proability distributions of *A*_1_ and *A*_2_ for *N* = 1000 and *ρ* = 0.5 estimated by maximum likelihood (ML) and Markov-Chain Monte-Carlo (MCMC). The results of the simulations are summarized in Table I. In terms of uncertainties we find that MCMC correctly estimates parameter uncertainties consistent with the distribution over a large sample, but that the maximum likelihood method underestimates the uncertainty by a constant factor that depends on the degree of correlation *ρ* or *C*. In Fig. 4 we show using a scatter plot that the individual uncertainties of MCMC and ML are correlated. This fact is important since MCMC is much more computationally expensive. For large datasets, we can use the maximum likelihood method to run over the whole dataset, and then we can use MCMC on a few samples to determine the factor by which the ML method underestimates the uncertainties. In our example, maximum likelihood underestimates the uncertainties by a factor of 1.5-2 over a wide range of *ρ*. We believe this underestimation results from the fact that the distributions in Fig. 3 are not Gaussian. The ML method assumes that all posterior distributions are Gaussian and that the width can be determined by the second derivative of the log likelihood. This is clearly not the case. The fact that we underestimate the uncertainties indicates that the curvature of our distribution at the maximum is greater than a corresponding Gaussian distribution.

**FIG. 3:**
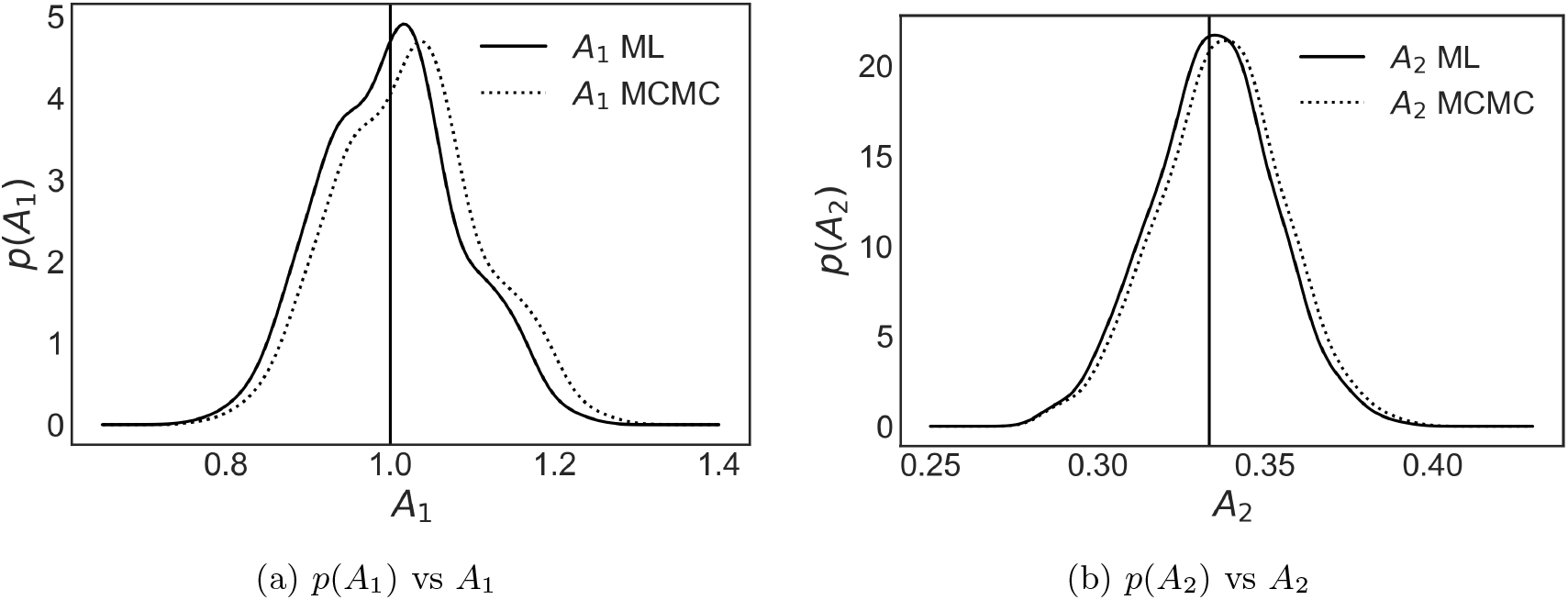
Comparison of MCMC and maximum likelihood for *ρ* = 0.5 and *D* = 1*, A*_1_ = 1, ∆*t* = 0.3*, N* = 1000. The vertical bar indicates the true expected value

**TABLE I:**
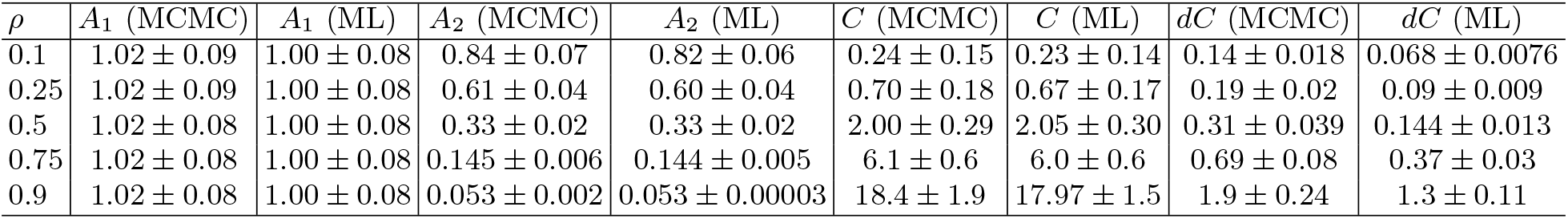
Comparison of parameter estimates using maximum likelihood and Markov-Chain Monte-Carlo methods for different *ρ* at *D* = 1*, A*_1_ = 1, ∆*t* = 0.3*, N* = 1000

**FIG. 4:**
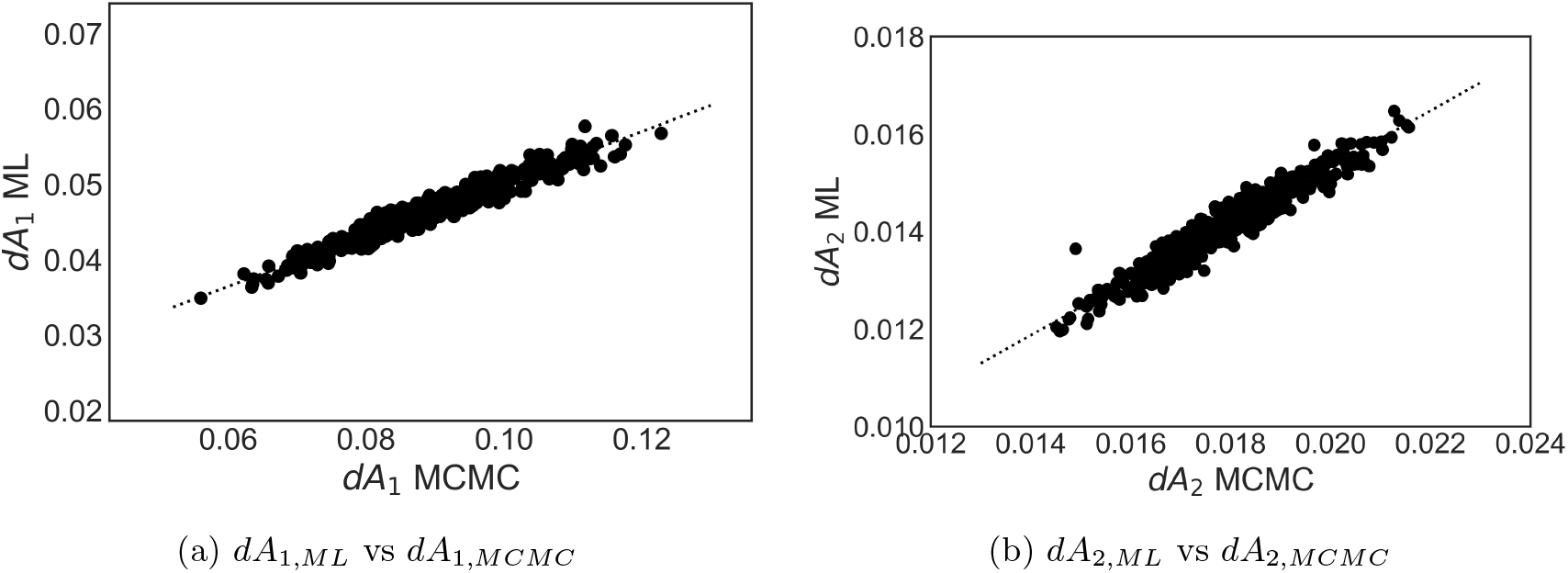
Correlation between the uncertainties of A estimated by maximum likelihood and MCMC for *ρ* = 0.5 and *D* = 1*, A*_1_ = 1, ∆*t* = 0.3*, N* = 1000. The dotted line represents a linear fit. For this *ρ* the maximum likelihood underestimates the errors by a factor of 2

## E. Application to resting-state functional Magnetic Resonance Imaging

To illustrate the power of our correlation method, we analyzed functional Magnetic Resonance Imaging (fMRI) time-series data from control 7T-fMRI resting-state experiments that were taken from [13]. In particular, we extracted neural activation time-series from four brain regions that make up the default mode network [14] (LLP = left lateral parietal cortex, RLP = right lateral parietal cortex, MPFC = medial prefrontal cortex, PCC = posterior cingulate cortex) that activates when individuals are focused on their internal mental-state processes, such as self-referential processing, interoception etc. and not performing a specific task. Fig. 5 shows the time series of a single subject. Traditially, correlations between brain regions are evaluated using Pearson correlation coefficients. To ensure that our coupled OU model is applicable, we determined the skewness and kurtosis of all time series to show that they are Gaussian. None of the distributions had an absolute value of skewness/kurtosis of more than 0.5, which is considered safe for applying Gaussian statistics (see for example [15]). Next, we have to show that the time series exhibit an exponentially decaying autocorrelation function. For this, we applied our maximum likelihood method for OU processes to determine the relaxation times for all time series. We found that *τ_MPFC_* = 4.8±0.7*sec*, *τ_PCC_* = 4.8±0.7*sec*, *τ_RLP_* = 4.1±0.5*sec*, *τ_LLP_* = 3.8±0.5*sec*. The decay-times of all brain regions are in within one or two standard deviations of each other, justifying the assumption made in eq. 3. Fig. 6 shows the coupling coefficients of five subject estimated by our method. Here we show the estimates from MCMC, but the same mean values are obtained by using the maximum likelihood method with uncertainties that are about a factor of 3 smaller than the MCMC estimates (see the Discussion in Simulation section). We chose four of six possible pairings of brain regions to illustrate the additional information that our method provides over Pearson correlations. By obtaining not only the strength of coupling but also the uncertainties of the value given the data, we can distingish between variance due to measurement and variance due to difference between subjects. Panel (a) illustrates a case where the uncertainty of the measurements is smaller than the variance of the mean. In this case this is due to an outlier. Here the traditional analysis would have sufficient to detect an outlier. In Panel (b) and (d) the variance of the mean is in between the variances of the individual datapoints and because we know each measurements uncertainty, we can determine how much of the variance of the mean is due to the variance between subjects and due to measurement error. Panel (c) illustrates a case where the uncertainty of the measurement is larger than variance of the mean.

**FIG. 5:**
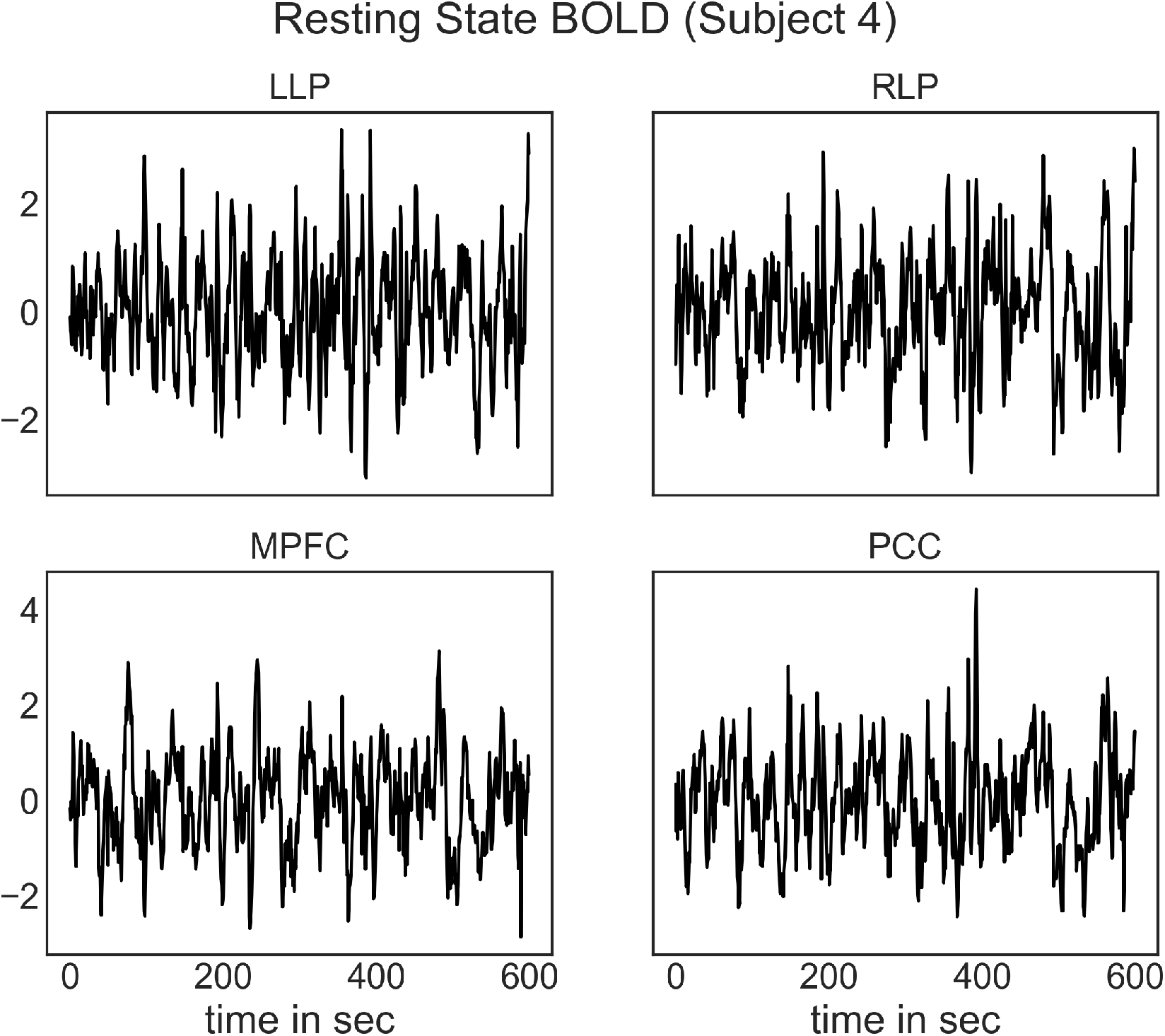
Time series of fMRI-BOLD signals of four brain regions that constitute the default mode network (LLP = left lateral parietal cortex, RLP = right lateral parietal cortex, MPFC = medial prefrontal cortex, PCC = posterior cingulate cortex)

**FIG. 6:**
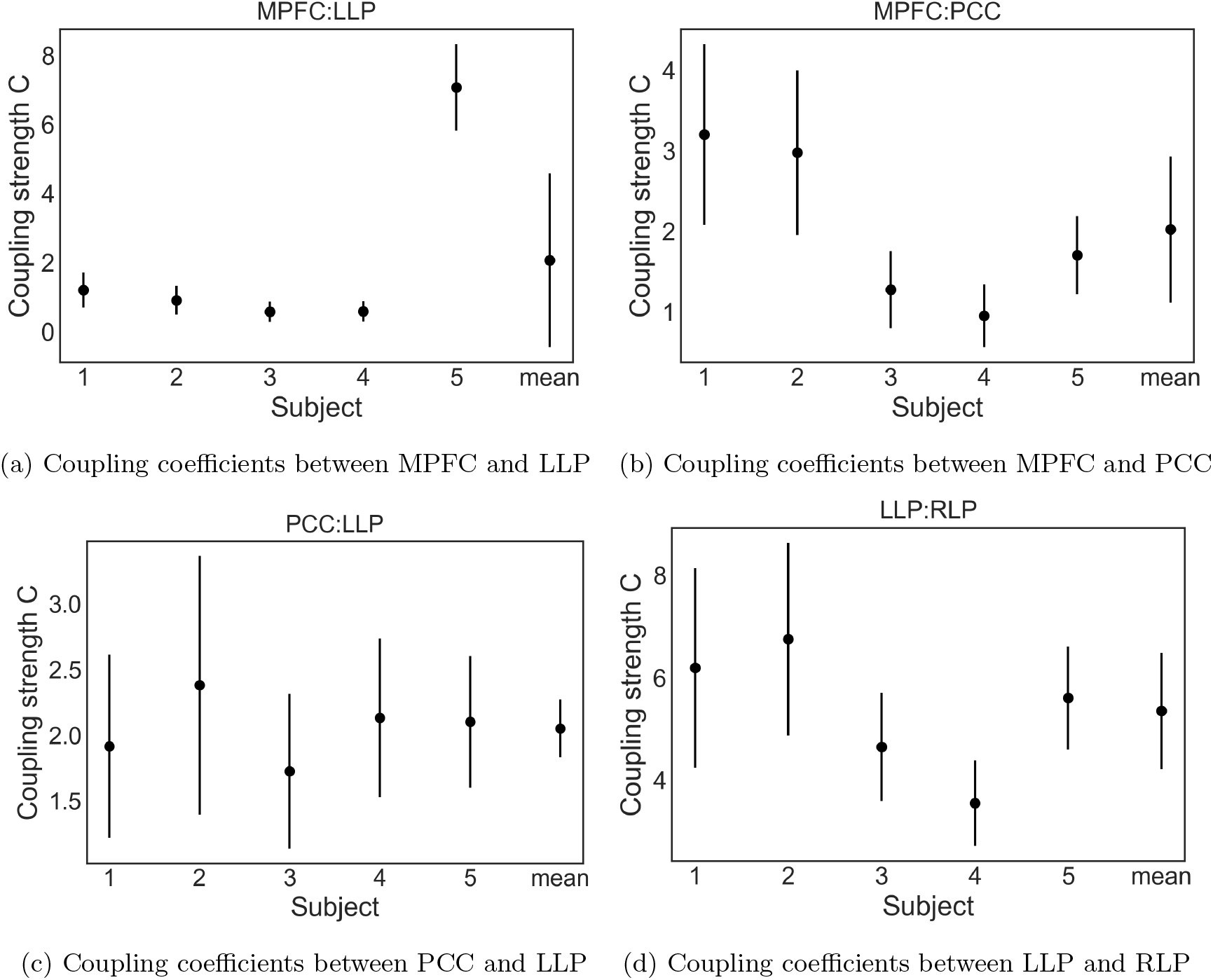
Coupling coefficients between default mode network brain regions across several subjects and the mean and standard deviation between subjects. For comparison to Pearson correlation coefficients we can use *ρ* = *C/*(2 + *C*) (*ρ* = 1*/*3 for *C* = 1, *ρ* = 0.6 for *C* = 3, and *ρ* = 0.75 for *C* = 6). The coupling coefficients and their uncertainties were determined by MCMC. We obtained the same mean coupling coefficients using the maximum likelihood method but with uncertainties that are about a factor of 3 smaller than the MCMC estimate.

## F. Summary

In the previous sections we developed and validated a maximum likelihood method to estimate the correlation between time series with an intrinsic relaxation time (OU process). As compared to Pearson correlation, our method not only estimates the correlation parameter, but also its confidence interval, which in our opinion is the most useful feature of our method. Knowing the confidence interval of individual measurements allows us to distingish between measurement error (confidence in a single measurement) and the variance that results from the variance of the sample. In the biological sciences, we often find large biological variance that is manifested in cell-cell variability of genetically identical cells or in the variance in the measurement of brain circuitry between (different subjects) but also within subjects (same subject at different days). In order to characterize these biological variances, biophysical methods have to be developed that are capable of measuring quantities accurately enough as to distingish between measurement error and biological distributions. Only if the individual error of a measurement is known we can attempt such a deconvolution. We hope that our method will help in achieving this goal.

## ACKNOWLEDGMENTS

We wish to acknowledge funding by the the National Institute of Drug Abuse (SBIR Phase 1 & Phase 2 Grant 1R44 DA043277-01) and the National Heart, Lung and Blood Institute (Grant 5U01HL12752202)

